# Nanoblades allow high-level genome editing in organoids

**DOI:** 10.1101/2022.08.04.502859

**Authors:** Victor Tirolle, Adrien Krug, Emma Bokobza, Mattijs Bulcaen, Marjolein M. Ensinck, Maarten H. Geurts, Delilah Hendriks, François Vermeulen, Frédéric Larbret, Alejandra Gutierrez-Guerrero, Louise Medaer, Rik Gijsbers, Philippe E. Mangeot, Hans Clevers, Marianne S. Carlon, Frederic Bost, Els Verhoeyen

## Abstract

Genome engineering has become more accessible thanks to the RNA programmable endonucleases such as the CRISPR/Cas9 system. However, using this editing technology in synthetic organs called ‘organoids’ is still very inefficient. This is due to the delivery methods used for the CRISPR-Cas9 machinery, which include electroporation of CRISPR/Cas9 DNA, mRNA or ribonucleoproteins (RNPs) containing the CAS9-gRNA complex. However, these procedures are toxic to some extent for the organoids. Here we describe the use of the ‘Nanoblade’ technology, which outperformed by far knock-out (KO) levels achieved to date by gene editing in murine and human tissue derived organoids. We reached up to 80% of gene KO in organoids after treatment with nanoblades. Indeed, high-level nanoblade-mediated KO for the androgen receptor (AR) encoding gene and the cystic fibrosis transmembrane conductance regulator (CFTR) gene was achieved with single gRNA or dual gRNA containing nanoblades in murine prostate and colon organoids. Likewise, nanoblades achieved high levels of gene editing in human organoids ranging between 20% and 50%.

Most importantly, in contrast to other gene editing methods, this was obtained without toxicity for the organoids. Only four weeks are required to obtain stable gene KO in organoids and nanoblades simplify and allow rapid genome editing in organoids with little to no side-effects such as possible unwanted INDELS in off-target sites.

## INTRODUCTION

Organoids are a self-organized three-dimensional (3D) culture system derived from embryonic, adult or induced pluripotent stem cells (ESCs, ASCs and IPSCs, respectively) (Sato et al., 2011b, 2011a, 2009). They recapitulate architecture, composition and functionality of their original epithelial tissues more faithfully than the traditionally used two-dimensional immortalized cell lines (Clevers, 2016; Drost et al., 2016). This model can be used to study stem cell differentiation and spatial organization processes (Matano et al., 2015). Thus, organoid technology is ideal for deciphering the role of genes involved in organogenesis or human pathologies (Drost et al., 2017; Dutta et al., 2017). Therefore, efficient approaches to edit the genome of mouse tissue-derived organoids and organoids derived from human adult stem cells (ASCs) are urgently required (Artegiani et al., 2020, 2019; Fujii et al., 2015; Geurts et al., 2021a; Hofer and Lutolf, 2021; Schwank et al., 2013).

Gene editing consists in manipulating the genome to induce gene silencing, gene modification or transgene integration at a precise locus (Doudna and Charpentier, 2014). In contrast to ectopic DNA sequence insertion using integrative vectors, genome editing allows more physiological gene manipulation. Precise genome editing also avoids gene silencing of non-targeted genes and adverse mutagenic effects such as insertional mutagenesis. Gene editing is based on the induction of double strand breaks (DSBs) at a precise locus (Gaj et al., 2013). It relies on engineered nucleases with a sequence specific genomic DNA-binding domain coupled to a non-specific endonuclease. Several engineered nucleases have been developed, such as zinc finger nucleases (ZFN) and transcription activator-like effector nucleases (TALENs). More recently, the clustered regularly interspaced short palindromic repeats (CRISPR)/associated protein 9 (Cas9) technology has been introduced in the field (Doudna and Charpentier, 2014). This relies on an endonuclease that uses a single guide RNA sequence (sgRNA) to introduce a site-specific DSB in the targeted DNA. The most common repair that occurs after a DSB is non-homologous end-joining (NHEJ). This consists in the fusion of the two DNA ends and can lead to the insertion or deletion of a few base pairs (INDELs). The frame shifts induced by these INDELs will modify partially or totally gene transcription and translation. Alternatively, homology directed repair (HDR) can occur when a donor sequence, with locus-specific homology arms, is available in addition to the endonuclease. This allows the introduction of a specific DNA alteration, like single-base substitution and also insertion of site-specific ectopic DNA sequences. However, HDR is restricted to the S/G2 phases of the cell cycle, and generally less efficient as compared to NHEJ, which makes this approach challenging for scientists in some primary cell models (Hustedt and Durocher, 2017).

The CRISPR/Cas9 technology has revolutionized the methodology to induce gene-specific knock-outs due to its high specificity, easy design and high efficiency in genetic manipulation of cell lines and primary cells. Endonucleases and sgRNAs can be introduced in the cell, by using CRISPR/Cas9 and sgRNA encoding retroviral vectors (Heckl et al., 2014). Alternatively, electroporation of plasmids or mRNA encoding the Cas9 have also been used (Hendel et al., 2015; Mandal et al., 2014). The main approach used in ASC-derived organoids is still electroporation of plasmids coding for the Cas9 endonuclease and the sgRNA (Fujii et al., 2015). Despite the fact that electroporation is widely used today, this technique leads to low efficiency and induces a high level of cell death (Artegiani et al., 2020; Fujii et al., 2015). Nevertheless, a recent study reported that the electroporation of ribonucleoprotein (RNP), combining Cas9 protein and a synthetic single-strand gRNA, provided higher gene editing efficiency into cells (Dawei et al., 2020).

Since KO efficiency in organoids is low, knock-in technologies have been developed to identify these KO events. The method uses a donor template coding for an expression cassette driving a fluorescent protein such as GFP flanked by homologous genome sequences (homology arms). Through recombination at the DSB induced by Cas9, this template permitted easy identification of KO/knock-in organoids through GFP expression (Okamoto et al., 2021). Alternatively, a NHEJ strategy relying on the piggybac-transposase system to integrate hygromycin/GFP was used to screen for fluorescence or drug resistance (Artegiani et al., 2020).

Gene editing has already been performed in several types of organoids such as rectal organoids. Cystic fibrosis transmembrane conductance regulator (CFTR) is an integral membrane protein that forms an anion channel activated by cAMP-dependent phosphorylation (Moran, 2010). An assay to reveal CFTR function used Forskolin induced fluid secretion into the organoid lumen resulting in swelling of rectal organoids (Dekkers et al., 2013). A study published in 2013 reported the use of the CRISPR/Cas9 genome editing system to correct the *CFTR* gene and thus restoring forskolin-induced swelling in patient-derived intestinal rectal organoids (Schwank et al., 2013).

Androgenic signals through the androgen receptor (AR) are required for luminal differentiation of some prostate basal stem cells. AR deletion in luminal cells has been shown to alter cell morphology and induce transient overproliferation (Xie et al., 2017). The first study that established prostate organoid cultures (Karthaus et al., 2014), showed that dihydrotestosterone (DHT) deprivation of CD26 + luminal cell-derived organoids disrupted lumen formation. Therefore, *AR*-KO prostate organoids generated by gene editing are expected to be compact even when stimulated with DHT. Therefore, both *CFTR* and *AR* are model target genes to evaluate gene editing in organoids.

Recently, a gene editing tool called nanoblades, based on a virus like particle (VLP) derived from a murine leukemia virus (MLV) has been developed (Bernadin et al., 2019; Gutierrez-Guerrero et al., 2021, 2021; Mangeot et al., 2019a). This technology uses the VLPs to introduce the Cas9/sgRNA RNPs into cells. These nanoblades carry the Cas9 proteins complexed with the sgRNAs, and are devoid of a viral genome, which allows a very quick and transient delivery of the gene editing machinery into the targeted cells. Here, we demonstrated that nanoblades allowed a very high KO efficiency for *AR and CFTR* in murine prostate and both murine and human rectal organoids, respectively. Moreover, this KO efficiency was accompanied by low toxicity and no obvious off-target effects. This facilitates the generation of gene KO in organoids since it does not require the use of a reporter encoding knock-in cassette to facilitate KO detection.

## MATERIALS AND METHODS

### Organoid culture

Mouse prostate organoids were generated as described by Drost *et al*. (Drost et al., 2016). Briefly, the prostates of 7-8-week-old male C57BL/6JOlaHsd mice (ENVIGO, Cannat, France) were isolated, and minced in small fragments. Tissue fragments were digested with type II collagenase at 5mg/ml for 1 hour at 37°C and subsequently incubated with 1X TrypLE (Gibco, Waltham, USA) for 15 minutes at 37°C. Cells were washed between each step with 15 ml of advanced complete DMEM/F12 (adDMEM/F12 containing penicillin/streptomycin, 10 mM HEPES and 2 mM GlutaMAX (100× diluted)) (ThermoFisher, Waltham, USA). Then cells were passed through a 70-μm cell strainer (ThermoFisher, Waltham, USA) to remove large cell debris. The cells were then labeled with CD49f-APC (Thermofisher, Waltham, USA, clone GoH3; RRID: AB_891474) and CD24-PE (Biolegend, clone M1/69, RRID: AB_493485) conjugated antibodies and were sorted for the CD49f+ CD24-cells by flow cytometry (BD FACSAria™ Cell Sorter). The isolated cells were then washed with complete adDMEM/F12 and after centrifugation at 200g for 5 minutes, the cell pellet was re-suspended in growth factor reduced phenol red-free matrigel (Corning, NY, USA) at a concentration of 200 cells per microliter of matrigel. Mouse prostate organoids were maintained in complete adDMEM/F12 supplemented with 50 X diluted B27 (Life Technologies Carlsbad, USA), 1,25 mM N-acetyl-l-cysteine (Sigma-Aldrich, Saint-Louis, USA), 50 ng/ml EGF (PeproTech, Cranbury, USA), 200 nM A83-01 (Tocris Bioscience, Bristol, UK), 1% Noggin conditioned medium, 10% R-spondin 1 conditioned medium, 1 nM DiHydro-Testosterone (Sigma-Aldrich, Saint-Louis, USA) and 10 μM Y-27632 dihydrochloride (PeproTech Cranbury, USA). Culture medium was replenished every two days.

Mouse colon organoids were established from colon of male C57BL/6JOlaHsd mice (ENVIGO, Cannat, France) and maintained in complete adDMEM/F12 supplemented with 50x diluted B27 (Life Technologies, Carlsbad, USA), 100x diluted N2 (Life Technologies Carlsbad, USA), 1 mM N-acetyl-l-cysteine (Sigma-Aldrich, Saint-Louis, USA), 50 ng/ml EGF (PeproTech Cranbury, USA), 1% Noggin conditioned medium, 20% R-spondin 1 conditioned medium,50% Wnt3a conditioned medium and 10 μM Y-27632 dihydrochloride (PeproTech Cranbury, USA). Culture medium was replenished every two days.

For passage of the organoid cultures, each drop of matrigel containing the organoids was resuspended in 1 ml of ice cold complete adDMEM/F12, which were centrifuged for 5 minutes at 200g at 4 ° C. Prostate and colon organoids were then incubated in TrypLE for 5 and 3 minutes, respectively, at 37 ° C and mechanically dissociated by pipetting up and down with a P10 / P1000 pipet tip. Cells were then washed with complete adDMEM/F12 and centrifuged. Cells were seeded back in matrigel at a concentration of 100 cells/μL. The appropriate medium was added to the cells and replenished every two days. Organoids were passaged every 7 to 10 days.

Human rectal organoids were generated and cultured as previously described (Vonk et al., 2020). Informed consent was obtained prior to biopsy collection, in accordance with the ethical committee of UZ Leuven (S56329). Human rectal organoids were generated form rectum biopsies of PwCF as previously described and biobanked after successful culture (Vonk et al., 2020). Organoids from the biobank were tawed and cultured for evaluation in this study.

### Plasmids

All the plasmids (Gagpol MLV, GagMLV-Cas9, VSV-G) and gRNA expressing plasmids to produce the nanoblades were described previously (Gutierrez-Guerrero et al., 2021; Mangeot et al., 2019a) and are available at Addgene (https://www.addgene.org). The BaEVRless envelope glycoproteins were previously described (Girard-Gagnepain et al., 2014a) and can be obtained under material and transfer agreement (MTA) from the corresponding author E. Verhoeyen. Both envelope glycoproteins (VSV-G and BAEV) were expressed in the phCMV-G expression plasmid (Maurice et al., 2002). The HIV-SFFV-eGFP vector and the HIV CMV-eGFP lentiviral vector encoding plasmids to produce the GFP murine and human organoid cell line were described previously (Levy et al., 2016; Vidović et al., 2016).

### Lentiviral production

Lentiviral vectors encoding the reporter gene eGFP were produced as described previously (Girard-Gagnepain et al., 2014a).

### sgRNA design

The sgRNAs of the different targeted genes were cloned into the gRNA expressing plasmid using the BsmBI restriction site (Mangeot et al., 2019b). The sequences of the sgRNA are as follows:

GFP_B182 5’-CGAGGAGCTGTTCACCGGGG-3’

Ms_AR_1 5’-ATGTACGCGTCGCTCCTGGG-3’

Ms_AR_2 5’-TTCAAGGGAGGTTACGCCAA-3’

Ms_CFTR_1 5’-GTGGCGATCATGTTGCTGCG-3’

Ms_CFTR_2 5’-AGTTCCGGATTCTGAACAGG-3’

Hu_CFTR 5’-GGAGAACTGGAGCCTTCAGA-3’

### Production of Nanoblades

Nanoblades were produced in human HEK293T kidney cell line (ATCC CRL-3216). 5×10^6^ cells were seeded per 10 cm culture plate 24h prior to transfection. To produce nanoblades, plasmids coding for MLVGagPol (3 μg), GagCas9 (3 μg), BaEVRless (2 μg), VSVG (2 μg) and gRNA expression plasmid (6 μg) were co-transfected with transfection reagent at a 2.6/1 ratio of Polyethylenimine (PEI; Polysciences, Warrington, USA) (μg)/total DNA (μg). The medium was replaced 24 hours after transfection by 6 ml of OptiMEM (Life Technologies Carlsbad, USA) supplemented with penicillin – streptomycin (1%) and HEPES (1%). The next day, Nanoblade-containing medium was clarified by centrifugation (2000 rpm for 5 min) and filtered through a 0.45 μm pore-size filter before centrifugation overnight (3000g for 16h at 4 °C). The supernatant was carefully removed by aspiration to obtain 100-150x concentration of the nanoblade preparation. Nanoblades can be stored at -80 °C.

### Cas9 quantification in the nanoblades by ELISA

Recombinant Cas9 (New England Biolabs, USA) was used to generate a standard curve (20µM, 6 serial dilutions of 1/2), while the nanoblade supernatants were diluted 1/200 and 1/400. The dilutions were performed in coating buffer (1% Triton) and were then coated onto 96-well-plates by incubation overnight at 4°C. The following day, the wells were incubated with washing buffer (PBS/0.05% Tween) and blocked with PBS/0.05%Tween/3%BSA (Sigma). Subsequently the wells were washed and the primary anti-Cas9 antibody (Cas9-7A9-3A3, 14697P; Cell signaling Technology, Inc, USA) was added at 1/1000 dilution in PBS/3%BSA, and incubated at RT for 1 h, while shaking. Before and after 1 h incubation with a secondary anti-mouse HRP (F6009-X639F South biotech, USA) diluted 1/10000 in PBS/3%BSA, a wash-step was performed. Finally, the mixed TMB substrate solution, containing HRP substrate was added for 20 min (Bethyl, Inc Texas, USA). Stop reaction was added in each well and protein was measured at 450 nm in a Multiskan FC (Thermo Scientific).

### Nanoblade transduction procedure

#### Murine organoids

Organoids were resuspended in ice-cold complete adDMEMF/12, then centrifuged at 200g for 5 minutes at 4 °C. Once the supernatant was aspirated, 1X TrypLE was added to the tube and incubated for 5 minutes at 37 ° C while pipetting up and down to dissociate the organoids.

For mouse prostate and colon organoids, after washing the cells with complete adDMEMF/12, 1×10^4^ cells were distributed in a 96-well plate in 100 μl of complete adDMEMF/12 with 10 μM Y-27632, to which nanoblades (4 µmoles of Cas9 protein) were added. The plate was then centrifuged at 200g for 60 min at room temperature and incubated for 6 hours at 37 °C. Colon organoid cells were only incubated with nanoblades for 3 hours. The cells were then collected in a 15 ml tube containing 5 ml of complete adDMEMF/12 and centrifuged at 200g for 5 min. The supernatant was removed and the cells were resuspended in 100 μl of matrigel and drops of 40 μl were deposited in 24-well plates. After an incubation of 15 minutes at 37 ° C, 500 μl of mouse prostate organoid medium was added to each well as previously described. Alternatively, the 96-well plate can be coated with RetroNectin^®^ (Clontech/Takara; 12µg/ml PBS according to manufacturer’s recommendations) overnight at 37 °C before seeding the cells and addition of the nanoblades. Polybrene (6 μg/ml) was added in some conditions as indicated to the nanoblades to enhance the transduction efficiency.

#### Human organoids

Human intestinal organoids were trypzinized to single cell suspension and then human rectal organoids were centrifuged for 5 min at 200g. The cell pellet was resuspended with 10 μl of nanoblade preparation per 6×10^3^ cells (*CFTR* KO experiment) or per 1.4×10^4^ cells (eGFP KO experiment) and incubated at room temperature for only 10 minutes. The cells loaded with nanoblades were then resuspended with matrigel and seeded into a 24-well plate. The plate was incubated 15 min at 37°C. Subsequently, the complete organoid medium (Dekkers et al., 2013), supplemented with rho-kinase inhibitor Y-27632 (10 μM) was added for 3 days to the solidified matrigel drops.

Establishment of murine and human organoid lines expressing eGFP: we used a lentiviral vector (plasmid pHIV-SFFV-IRES-eGFP (Frecha et al., 2008), which contains an eGFP under the control of an SFFV promoter or a lentiviral vector pHIV-CMV-eGFP with eGFP under the control of the CMV promoter (Vidović et al., 2016). The transduction procedure is the same as the one used for the nanoblades. We applied a low MOI of 0.3 to ensure that we obtained only one integrated copy of the eGFP expression cassette into the genome of the mouse organoids. We then sorted the cells by flow cytometry for eGFP+ cells (BD FACSAria™ Cell Sorter). We seeded the sorted murine organoid cells in matrigel at a concentration of 200 cells/μL. Then the corresponding media was added and replenished every two days.

#### Flow cytometry analysis

Murine organoids were harvested with ice cold complete adDMEMF/12. After centrifugation at 250g for 5 min at 4 °C, the supernatant was aspirated and discarded, then 1x TrypLE (Gibco, Waltham, USA) was added to each tube and incubated for 5 minutes at 37 °C. After vortexing the tubes, up and down pipetting was performed to dissociate the organoids into single cell suspension. The cells were washed with complete adDMEMF/12, and then analyzed by flow cytometry to determine eGFP expression levels. Human rectal organoids were dissociated with 0,25% Trypsin/EDTA, fixed with 4% paraformaldehyde for 15 min and resuspended in PBS. In HEK293T cells expressing 3HA-CFTR, CFTR plasma membrane (PM) density was measured in the presence and absence of anti-hCFTR nanoblades as described previously (Ensinck et al., 2020).

#### Immunofluorescent staining and imaging

For confocal imaging, organoids were observed with an Evos optical microscope. For immunofluorescence staining, organoids were harvested and washed once with ice cold complete adDMEMF/12. They were fixed for 1 hour in 4% PFA, permeabilized with 0.5% Triton X-100 for 30 min and then blocked with 2% Fetal Bovine Serum (FBS) in PBS for 1 hour at room temperature (RT). Then the organoids were incubated for 2 hours at RT with primary antibodies against cytokeratin 8 (Abcam, Cambridge, UK, DSHB, TROMA-I, AB_531826), cytokeratin 5 (AlexaFluor647 coupled antibody, Abcam, Cambridge, UK, RRID AB_2728796), and androgen receptor (Abcam, Cambridge, UK, AB_10865716). Then organoids were washed with PBS, followed by an incubation with secondary antibodies coupled to an AlexaFluor 488 (Goat anti-Rabbit IgG, Thermofisher, RRID: AB_143165) and AlexaFluor 594 (Goat anti-Rat IgG Thermofisher, RRID: AB_10561522) for 1h30 at RT. DAPI was used to stain nuclear DNA. Organoids were mounted and imaged with a confocal Nikon microscope. Images were processed with ImageJ.

#### Determination of nanoblade toxicity on organoid development

The number of murine organoids was determined using a counting cell at 9 days after incubation with nanoblades and reseeding the cells in the respective organoid cultures.

The number of human rectal organoids was determined by automated counting of organoids in 1 picture 14 days after incubation with nanoblades and reseeding the cells in the respective organoid cultures. Replicates were obtained from different wells in one experiment and 2 independent experiments were performed.

#### Detection of nanoblade mediated on-target gene editing efficiency

After incubation with nanoblades, murine organoids were grown for 9 days. Genomic DNA (gDNA) was extracted using a genomic DNA extraction kit (Macherey-Nagel). Primers were designed to amplify approximately 400-700 bp of the region targeted by the guide RNAs (***Supplementary Table 5***). 50 ng of gDNA was used for PCR amplification. The size of the PCR fragments was checked using an agarose gel. PCR fragments were then purified using a gel extraction kit (New England Biolabs) and subjected to sanger sequencing. Sequencing results were analyzed for the percentage of INDEL using ICE (https://ice.synthego.com/#/) or DECODR (https://decodr.org/).

#### Off-target genome editing detection

The potential off-target sites for the guide RNAs used in this study were determined using Cas-OFFinder (http://www.rgenome.net/cas-offinder/). Some of the off-target sites with 2 or 3 mismatches for each guide RNA were selected for further analysis. Note, only off-target sites with a PAM sequence (NGG) were tested here. PCR primers were designed to amplify a fragment of about 400 to 700 bp flanking the identified potential off-target sites (***Supplementary table 5***). Genomic DNA was isolated and the PCR and fragment isolation were performed as mentioned above. The PCR fragments were then sequenced by Sanger sequencing. Sequencing results were analyzed for the percentage of INDELs using ICE (https://ice.synthego.com/#/) and DECODR (https://decodr.org/).

#### Statistical Analysis

At least 3 independent biological replicates were performed using for each a different nanoblade preparation, except for the human organoids where 2 independent experiments were performed. The statistical tests applied are indicated in the figure legends.

## RESULTS

### Nanoblades allow efficient genome editing in organoid cells with low toxicity

As an easy read-out for nanoblade gene editing efficiency in organoids, we chose initially to knock-out eGFP in prostate organoids expressing eGFP. First, we established a mouse prostate organoid cell line expressing eGFP. Organoid cells were transduced with a lentiviral vector encoding eGFP driven by the SFFV promoter at low vector doses (MOI of 0.3) to ensure only one integrated copy of the eGFP expression cassette per organoid cell. The eGFP positive cells were sorted to establish the eGFP organoid cell line.

To assess eGFP KO, we selected a sgRNA targeting eGFP that previously resulted in 75% of eGFP KO in bone marrow-derived macrophages (Mangeot et al., 2019a). The nanoblade treatment is schematically represented in ***Figure 1A***. Organoids were dissociated and incubated with the nanoblades for only 6 hours. Of note, absence of growth factors for more than 6h induced cell death. Moreover, a brief incubation with nanoblades was expected to be sufficient since receptor mediated entry of the nanoblades is a fast process as previously shown for other primary cells (Gutierrez-Guerrero et al., 2021; Mangeot et al., 2019a). The cells were then rinsed to remove the nanoblades and seeded in matrigel. Without any prior selection for gene edited cells, the single cells were grown for 9 days into organoids. Fluorescence microscopy revealed a strong reduction of eGFP+ organoids (***Figure 1B***). Additionally, the bulk population of dissociated organoids was quantified by FACS for nanoblade mediated loss of eGFP expression (***Figure 1C***). We observed that increasing doses of anti-GFP nanoblades permitted to increase gene editing and resulted in up to 80% gene edited cells when the highest doses of nanoblades applied (***Figure 1C***).

**Figure 1.**
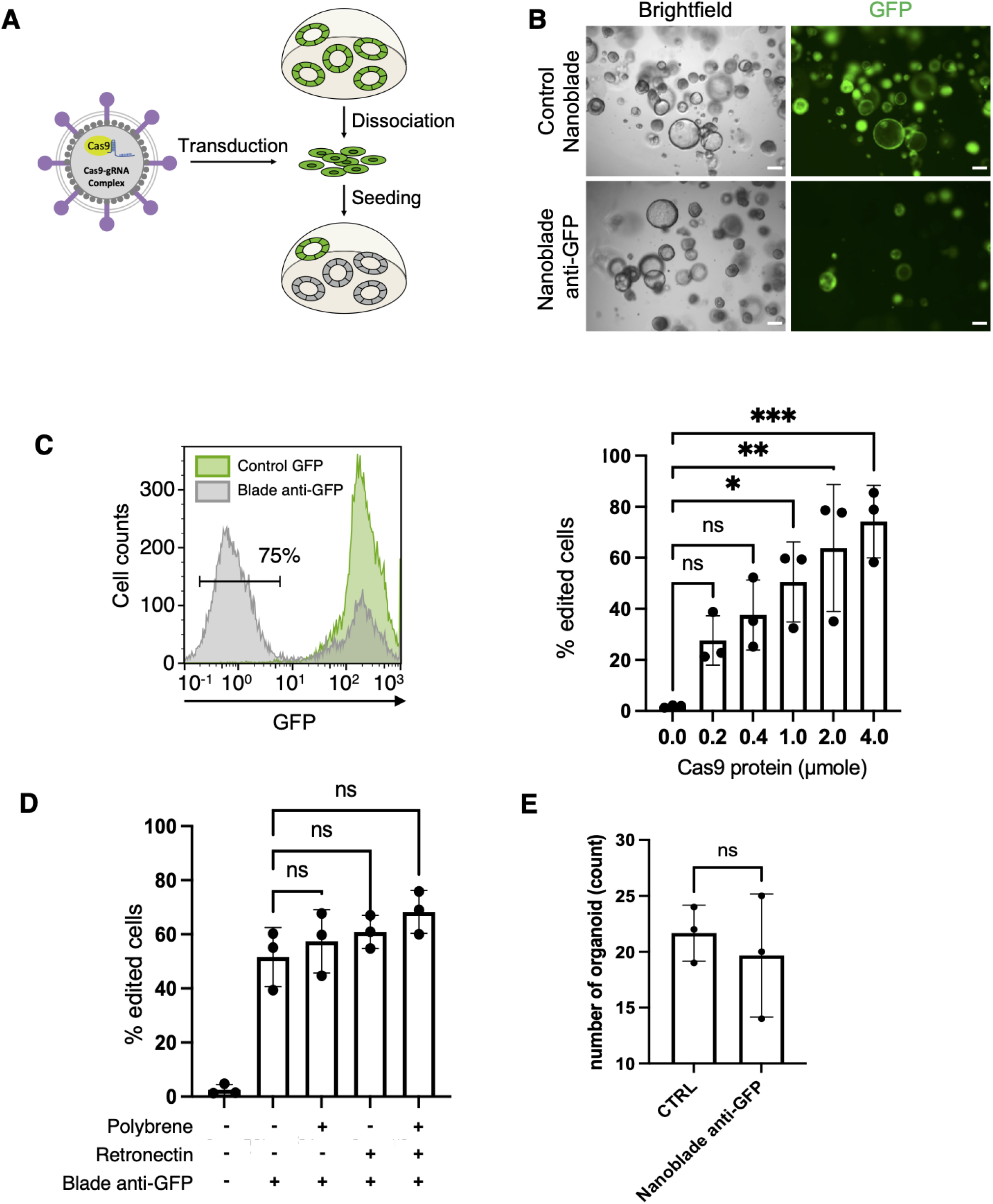
Strategy and efficiency of nanoblade induced knock-out in mouse prostate organoids. (**A**) Experimental workflow using nanoblades for gene editing of mouse prostate organoids. Organoids are dissociated, treated with the nanoblades, and seeded back into matrigel supplemented with growth factors inducing organoid generation. (**B**) Representative bright-field images of mouse prostate organoids 9 days after treatment with or without nanoblades targeting the eGFP-coding sequence. Scale bar, 200 µm. (**C**) Left panel, flow cytometry analysis of eGFP expressing organoids treated or not with nanoblades targeting the GFP-coding sequence. Right panel, flow cytometry analysis of eGFP expressing organoids treated with different quantities of anti-GFP nanoblades (means ± SD; n = 3, biologcial replicates; one-way ANOVA, Tukey’s multiple comparisons test; *p<0.05; * *p<0.01, ***p<0.001). (**D**) Flow-cytometry analysis of eGFP expressing organoids treated with fixed amounts of anti-GFP nanoblades with or without Polybrene and/or Retronectin (means ± SD; n = 3; biological replicates; one-way ANOVA, Tukey’s multiple comparisons test; ns= not significant). (**E**) The toxicity in organoids is evaluated, nine days after incubation of organoid cells with or without nanoblades by counting of the number of organoids developing per fixed number of cells seeded (means ± SD; n = 3, biological replicates; t-test, Mann Whitney test, non-parametric; ns= not significant).

In order to further improve the efficiency of nanoblades, we compared a constant dose of nanoblades in combination with two transduction facilitating agents, polybrene and/or retronectin (***Figure 1D***; (Girard-Gagnepain et al., 2014b)). No significant increase in gene editing efficiency using nanoblades was observed in the presence of retronectin or polybrene or both as compared to the control conditions in absence of these facilitating agents.

Of high importance, even after treatment of dissociated organoid cells with the maximum dose of nanoblades tested (4 µmoles of Cas9 protein) no significant toxicity was detected since the number of cells developing into organoids was equivalent in the presence or absence of nanoblade incubation (***Figure 1E***). These data suggested that nanoblades can reach high-level gene editing in organoids without toxic side-effects such as cellular toxicity.

### Nanoblades allow highly efficient gene knock-out in mouse prostate organoids

As a proof of concept, we chose to knock-out the gene coding for the androgen receptor (AR), a key protein in the development of mouse prostate and prostate organoids. Therefore, we designed two sgRNAs targeting exon 1 or exon 8 in the *AR* locus. The experimental set-up is shown in ***Figure 2A*** and as indicated can be performed in 4 to 6 weeks to obtain organoids KO for a specific gene. The layout of the experiment includes cloning of the gRNA in an RNA expression plasmid, followed by the nanoblade production carrying the *AR*-targeted sgRNAs. Further, dissociated mouse prostate wild type (WT) organoids were treated either with the nanoblades carrying the AR_1 sgRNA or the AR_2 sgRNA or both sgRNAs simultaneously. After 9 days of organoid culture, we revealed a strong phenotypic change in the organoids KO for *AR*, showing very compact round spheric structures without any lumen as compared to controls (***Figure 2B&C***). This is in agreement with the literature, since organoids that are not stimulated for the androgen pathway show a compact phenotype (Karthaus et al., 2014). Indeed, a cystic organoid phenotype is dependent on the stimulation of prostate luminal cells by dihydrotestosterone (DHT). Confocal microscopic acquisitions of the *AR* KO organoids showed absence of a lumen in the center of the organoid (***Figure 2C***). Labeling of AR with an anti-AR antibody also confirmed the absence of the AR protein in the compact *AR* KO organoids (***Figure 2C***).

**Figure 2.**
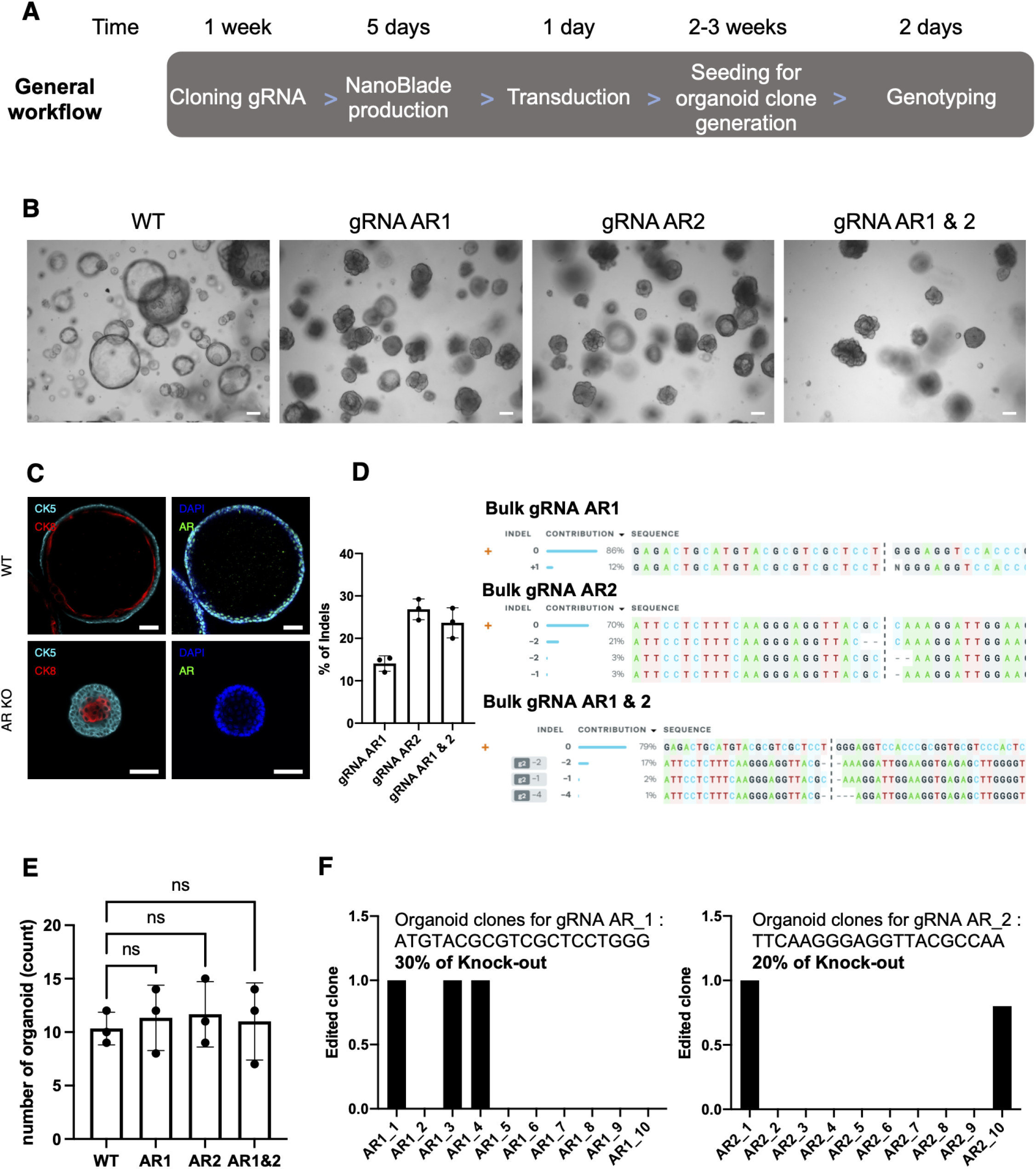
Nanoblades enable efficient generation of knock-out organoids from mouse prostate organoids. (**A**) Stepwise procedure and time line for nanoblade mediated gene KO in organoid cell lines (**B**) Bright-field images of mouse prostate organoids at 9 days after treatment with nanoblades targeting the Androgen-Receptor (AR). Two different sgRNAs (AR_1 and AR_2) were used to knock-out AR, either alone or in combination. Scale bar, 200 µm. ((Representative of n=3). (**C**) Immunofluorescence images of the center of WT and AR-KO mouse prostate organoids. Stainings were performed for the basal marker CK5 (cyan), luminal marker CK8 (red) and AR (green). Nuclei were stained with DAPI (blue). Scale bar, 50 µm. (Representative of n=3). (**D**) Bulk gDNA sequencing analysis of organoids treated with AR targeted nanoblades. Sanger sequencing was performed of the 557 bp PCR surrounding the AR target loci and was subjected to ICE analysis which revealed on-target INDEL frequencies at gRNA AR1 or AR2 target loci. Sequence decomposition of INDEL events is shown and are summarized in a histogram (means ± SD; n = 3; technical replicates). (**E**) The toxicity in organoids is evaluated nine days after treatment by counting the number of organoids treated or not with nanoblades targeting AR using a counting cell (means ± SD, n = 3, biological replicates, one-way ANOVA, Tukey’s multiple comparisons test; ns= not significant). (**F**) Single organoids were harvested, gDNA was isolated and sequenced for the loci targeted by the different AR guide RNAs. Sanger sequencing was performed of the 557 bp PCR surrounding the AR target loci and was subjected to ICE analysis which revealed on-target INDEL frequencies at gRNA AR1 or AR2 target loci.

Nine days after the treatment of the organoids with the nanoblades, we isolated genomic DNA (gDNA) of the bulk organoid population and amplified a 500 bp long sequence surrounding the genomic loci targeted by gRNA-AR_1 and gRNA-AR_2 using the AR_ctrl_fw and AR_ctrl_rv primers. Then, these PCR products were subjected to Sanger sequencing. Analysis of the resulting sequences by ICE confirmed that nanoblades incorporating only gRNA-AR_1 resulted in 12% INDELs, while gRNA-AR_2 containing nanoblades resulted in 27 % INDELs. When the two gRNAs AR_1 and AR_2 were combined in the same nanoblade, we reached 17% INDELs as shown by a representative ICE analysis, which were confirmed by DECODR analysis (***Figure 2D***). We also evaluated here the level of toxicity induced by treatment of the organoids with nanoblades targeting the *AR* gene (***Figure 2E***) by counting the number of outgrowing organoids nine days after the treatment with or without nanoblades. The *AR*-targeted nanoblades did not significantly affect survival of the cells nor their development into organoids, although the *AR* KO resulted in a strong change in phenotype.

Clonogencity is a major asset of the organoid model. Indeed, for prostate organoids, each reseeded progenitor cell will reform a new organoid. The organoid will then have the genetic identity of the initial KO cell. To confirm the efficiency of organoid KO line generation with nanoblades, we picked ten organoids per condition. For each of these organoids the *AR* locus targeted for gene editing was PCR amplified individually and evaluated for *AR* KO using ICE analysis. For the nanoblades containing gRNA AR_1, three out of ten organoid lines generated were KO for *AR*, which results in a level of 30% of *AR* KO organoids. For the nanoblades containing gRNA AR_2, two out of ten organoids picked and amplified were KO for *AR*, indicating 20% *AR* KO efficiency (***Figure 2F - figure supplement 1***).

### Nanoblades allow efficient gene editing in mouse colon organoids

Colon organoids are one of the first organoid models which have been developed (Sato et al., 2011a). They are also cultured in matrigel. The culture medium is slightly different in composition compared to that used for culture of mouse prostate organoids as described in materials and methods. The nanoblade-mediated editing efficiency and toxicity might be different according to the type of organoid; therefore, both parameters were evaluated in colon organoids.

As described above for prostate organoids, nanoblade gene editing efficiency in colon organoids was initially tested by knocking-out eGFP in colon organoids expressing eGFP. Similar to the prostate organoids, we established a mouse colon organoid cell line expressing eGFP. We incubated the dissociated colon organoid cells with anti-GFP nanoblades for 3 hours and grew them into organoids. Fluorescence microscopy demonstrated a strong reduction in eGFP+ organoids (***Figure 3A***). The bulk population of dissociated organoids was quantified by FACS for nanoblade mediated loss of eGFP (***Figure 3B***). We obtained with the colon organoids similar levels of KO as for the prostate organoids. Indeed, with the maximum dose of anti-GFP nanoblades we achieved gene editing efficiency of 70% for the colon organoid cells (***Figure 3C***). Equivalent to prostate organoids, after treatment of dissociated colon organoid cells with the maximum dose of nanoblades tested (4 µmoles of Cas9 protein) no significant toxicity was detected since the number of cells developing into organoids was equivalent in the presence or absence of nanoblades (***Figure 3D***).

**Figure 3.**
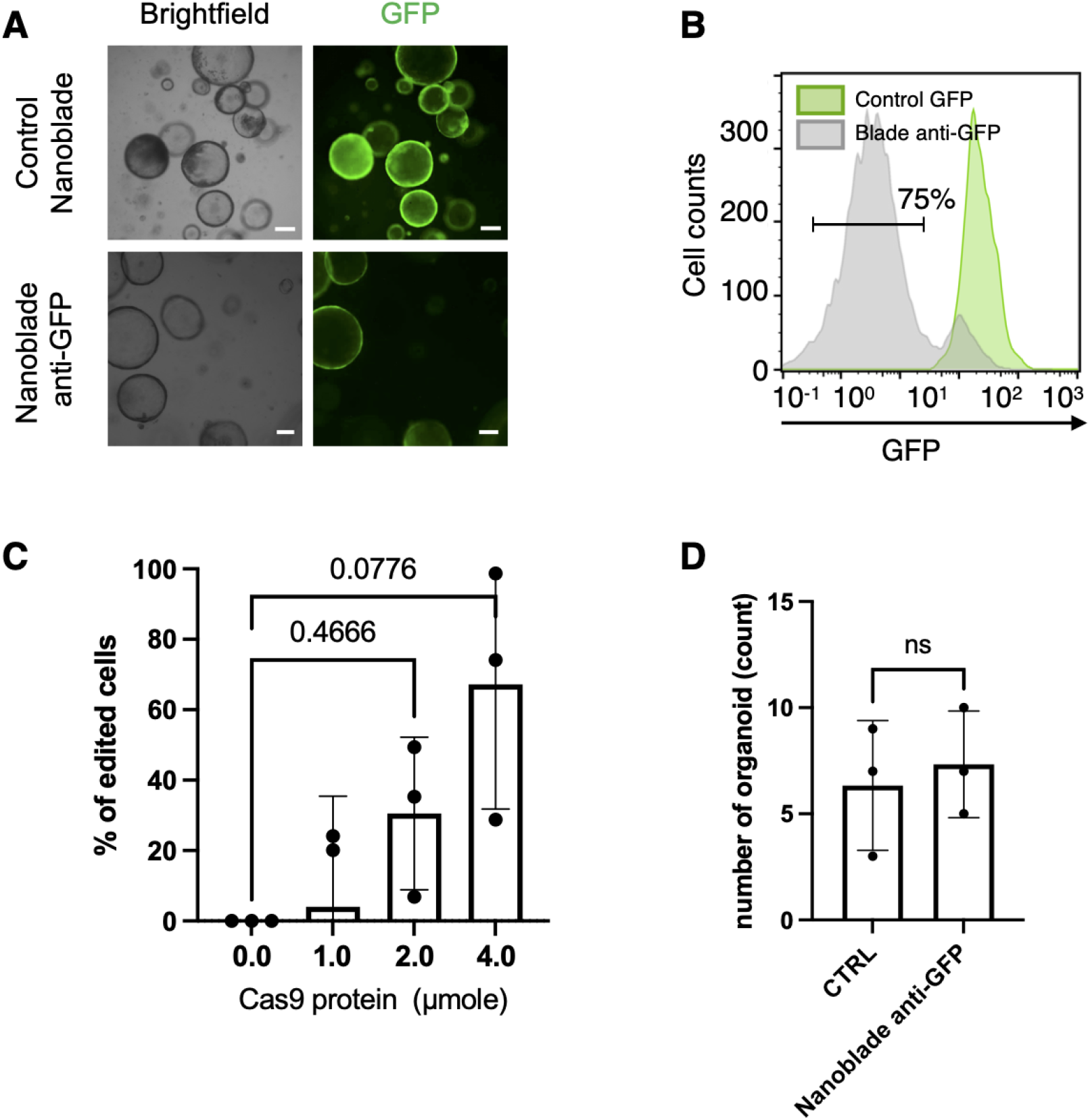
Efficient nanoblade induced knock-out in mouse colon organoids. (**A**) Representative bright-field images of mouse prostate organoids 9 days after treatment or not with anti-GFP nanoblades. Scale bar, 200 µm. (**B**) Flow cytometry analysis of eGFP expressing organoids treated or not with nanoblades targeting the eGFP-coding sequence. (**C**) Flow cytometry analysis of eGFP expressing organoids treated with different quantities of anti-GFP nanoblades (means ± SD; n = 3, biological replicates; one-way ANOVA, Tukey’s multiple comparisons test; ns = not significant). (**D**) Toxicity of nanoblades in organoids is evaluated nine days after treatment by counting the number of organoids treated or not with nanoblades using a counting cell (means ± SD; n = 3, biological replicates; t-test, Mann Whitney test, non-parametric; ns= not significant).

Overall, this first evaluation of nanoblade mediated gene editing in mouse colon-derived organoids shows a very high efficiency of genome editing with low toxicity, equivalent to what we observed for prostate organoids.

### Nanoblades allow efficient CFTR gene knock-out in mouse colon organoids

As mentioned previously, CFTR is an anion channel activated by cAMP-dependent phosphorylation (Moran, 2010). Previous studies have shown that CFTR is responsible for fluid secretion into the lumen and swelling of organoids (Dekkers et al., 2013; Geurts et al., 2021a; Schwank et al., 2013).

We used the same experimental approach as for *AR* KO in prostate organoids here for colon organoids (***Figure 2A***). We designed two gRNAs targeting the mouse *CFTR* gene (exon 14/exon27) and treated the colon organoids for 3 hours with nanoblades carrying either the CFTR_1, CFTR_2 or CFTR_1 & CFTR_2 gRNAs (***Figure 4A***).

**Figure 4.**
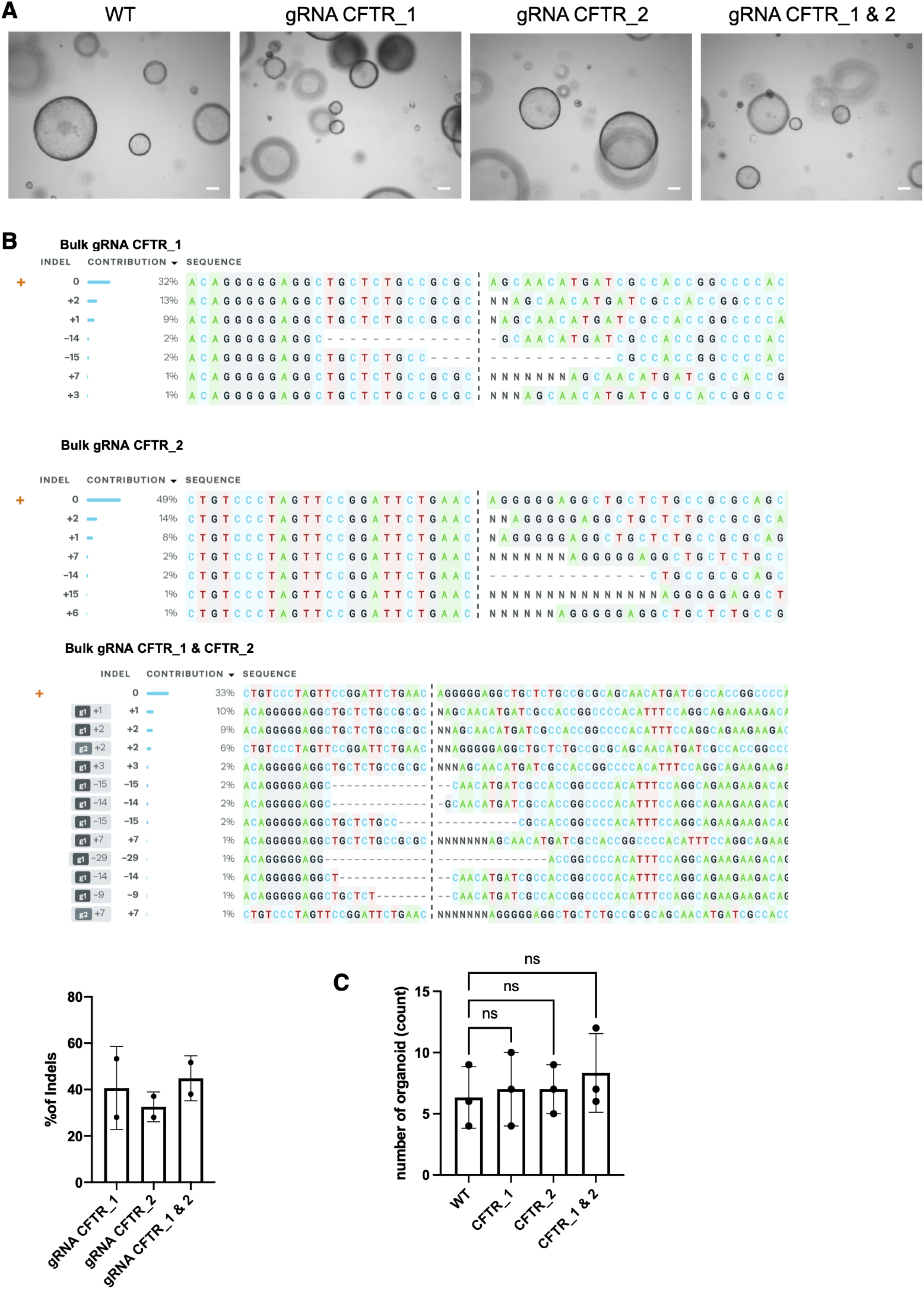
Nanoblades allow efficient knock-out of CFTR in mouse colon organoids. (**A**) Representative bright-field images of mouse prostate organoids 9 days after treatment with nanoblades targeting CFTR. Two different sgRNAs were used to knock-out CFTR, which were employed alone or in combination. (**B**) Bulk gDNA sequencing analysis of organoids treated with nanoblades targeting CFTR. Sanger sequencing was performed of PCR band including the CFTR target loci and was subjected to ICE analysis which revealed on-target INDEL frequencies at gRNA CFTR_1 or CFTR_2 target loci. Sequence decomposition of INDEL events is shown and are summarized in a histogram (means ± SD; n = 2, biological replicates). (**C**) Toxicity of nanoblades in organoids is evaluated nine days after treatment by counting the number of organoids treated or not with nanoblades using a counting cell (means ± SD; n = 3; biological replicates, one-way ANOVA, Tukey’s multiple comparisons test; ns= not significant).

Nine days after the treatment of the organoid cells with the nanoblades and their outgrowth in colon organoid medium, we isolated gDNA of the bulk organoid population (***Figure 4B***). We amplified a 625 bp genomic fragment around the loci targeted by the CFTR_1 sgRNAs or CFTR_2 sgRNA. The INDEL analysis performed by ICE software, showed an average INDEL frequency of 40% for CFTR_1 sgRNA, 32% for CFTR_2 sgRNA. The percentage of INDELs reached 44% when both guide RNAs were provided by the same nanoblade preparation.

Toxicity was also evaluated by counting the number of organoids that developed nine days after treatment in the presence or in the absence of the nanoblades (***Figure 4C***). Here again the nanoblades did not induce toxicity compared to the control conditions.

Nanoblade-mediated gene editing was thus also highly efficient for mouse colon organoids underlining the ease and versatility of nanoblades for gene editing and KO generation in the mouse organoid model.

### Nanoblades do not induce INDELs at the analysed off-target sites in murine organoids

Another advantage of the nanoblades is that we provide Cas9 as a protein. The CRISPR/Cas9 gRNA complexes, so-called RNPs, contained in the VLPs are only transiently present once delivered into the targeted cell (Gutierrez-Guerrero et al., 2021). The fact that Cas9 is present in the cell for a limited time should decrease the number of cuts in off-target sites as shown previously (Mangeot et al., 2019b). Using the Cas-OFFinder software, we predicted potential off-target sites for each of our sgRNAs (***Supplementary Tables 1 & 2***). We included in our analysis only the off-target sites containing a protospacer adjacent motif (PAM). We assessed two off-target sites for the AR_1 sgRNA with 3 mismatches since no off-target sites with 1 or 2 mismatches were identified by Cas-OFFinder (***Figure 5A - supplementary Table 1***). One was located on chromosome 9 and the other one on chromosome 13. We designed PCR primers (see material and methods) to amplify by PCR a 602 bp region around the chromosome 9 off-target site, and a 622 bp fragment including the potential chromosome 13 off-target site. For the AR_2 sgRNA off-target no sequences with 1, 2, or 3 mismatches were predicted and only off-target sites with 4 mismatches were identified compared to the target locus, of which we randomly choose 2 for off-target INDEL analysis (***Figure 5A - Supplementary table 2***).We amplified by PCR the off-target regions in the gDNA isolated from the organoid bulk treated with either AR_1 or AR_2 sgRNA and sanger sequenced the PCR products. Analysis for INDELs by ICE, showed that no gene editing was detected in the off-target sites analyzed (***Figure 5B***). This was confirmed by DECODR analysis.

**Figure 5.**
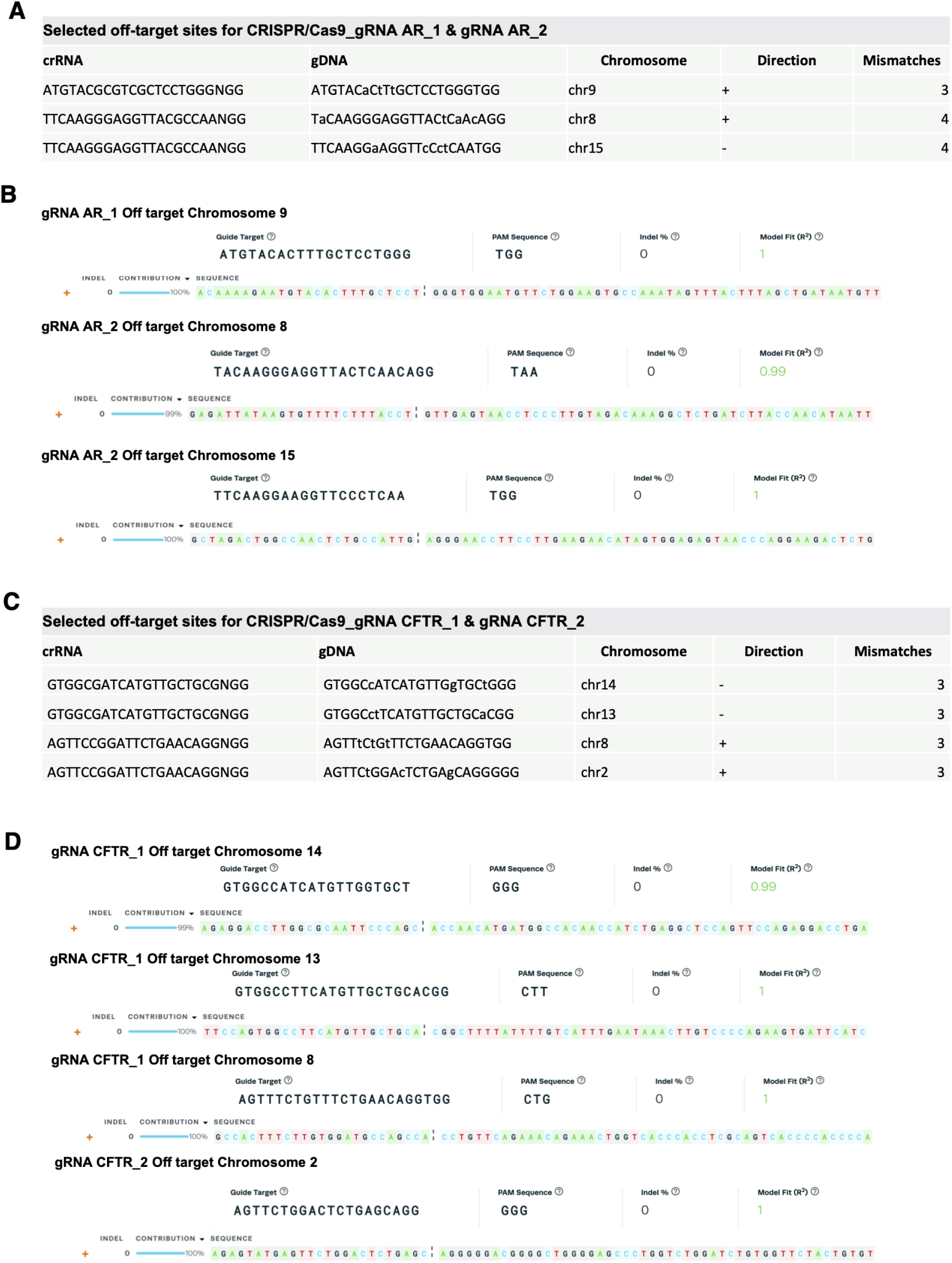
Off-target genotoxicity was not detected upon nanoblade-mediated KO of genes in organoids. (**A**) Cas-OFFinder algorithm was used to identify potential off-target sites for the sgRNAs AR1 and AR2 of which 3 are listed (see supplementary Table 1). (**B**) Sanger sequencing of sgRNAs AR1 and AR2 potential off-target loci subjected to ICE analysis revealed no off-target DSBs. (**C**) Cas-OFFinder algorithm was used to search for potential off-target sites for the sgRNAs CFTR_1 and CFTR_2 of which 4 are listed (see supplementary Table 1). (**D**) Sanger sequencing of sgRNAs CFTR_1 and CFTR_2 potential off-target loci subjected to ICE analysis revealed no off-target DSBs.

We also predicted the potential off-target sites for the sgRNAs CFTR_1 and CFTR_2 using the open access program Cas-OFFinder (***Supplementary Tables 3 & 4***; no more than 3 mismatches are listed). We proceeded as for *AR* gRNA off-targets and under these experimental conditions, no off-target gene editing was detected for the 4 different *CFTR* off-target positions analyzed (***Figure 5C&D***).

### Nanoblades allow efficient gene editing in human colon organoids

Human organoids such as intestinal or prostate organoids subjected to lentiviral transduction or electroporation to introduce the CRISPR/Cas9 machinery result in more cellular toxicity than their murine counterparts. Since the nanoblade technology performed gene editing in murine organoids without a significant toxic side effect, we wanted to evaluate the efficacy of nanoblade-mediated gene editing in human organoids. Firstly, we generated eGFP expressing human rectal organoids by short exposure to a LV expressing eGFP under the CMV promoter. Without sorting we exposed the human organoid cells (30% GFP+) to anti-GFP nanoblades (4 µmoles of Cas9 protein) for only 10 min to avoid toxicity on organoid growth since longer incubation to viral particles was toxic as shown previously (Vidović et al., 2016). The cells were then seeded in matrigel. Without any prior enrichment for edited cells, the cells were grown into organoids for 14 days. The bulk population of dissociated organoids was quantified by FACS for nanoblade-mediated loss of eGFP (***Figure 6A***). Over 40% of the GFP+ organoids lost eGFP expression and residual eGFP expressing cells showed reduced levels of eGFP (MFI), indicating that at least 40% gene editing in these human intestinal organoids. Subsequently, we designed a gRNA against human *CFTR*, which we first validated in the context of nanoblades (anti-hCFTR NB) in HEK293T cells. These HEK293T overexpress hCFTR due to an integrated *CFTR* cDNA copy tagged by an extracellular 3HA-Tag for detection by flow cytometry (Ensinck et al., 2020). After the treatment with the nanoblades, genomic DNA of the bulk HEK293T cells was isolated and the sequence surrounding the genomic locus targeted by the hCFTR sgRNA was amplified using the *hCFTR* Fw and Rv primers (***Supplementary Table 5***). These PCR products were then subjected to Sanger sequencing. Analysis of the resulting sequences by ICE confirmed that anti-hCFTR NBs resulted in over 90% INDELs (***Figure 6B***). The high-level KO of *hCFTR* in HEK293T was confirmed by almost complete disappearance of *hCFTR*-HA tagged surface expression (***Figure 6C***). As detected previously no cell death was induced by the NBs in 293T cells (***Figure 6D;*** (Gutierrez-Guerrero et al., 2021)).

**Figure 6:**
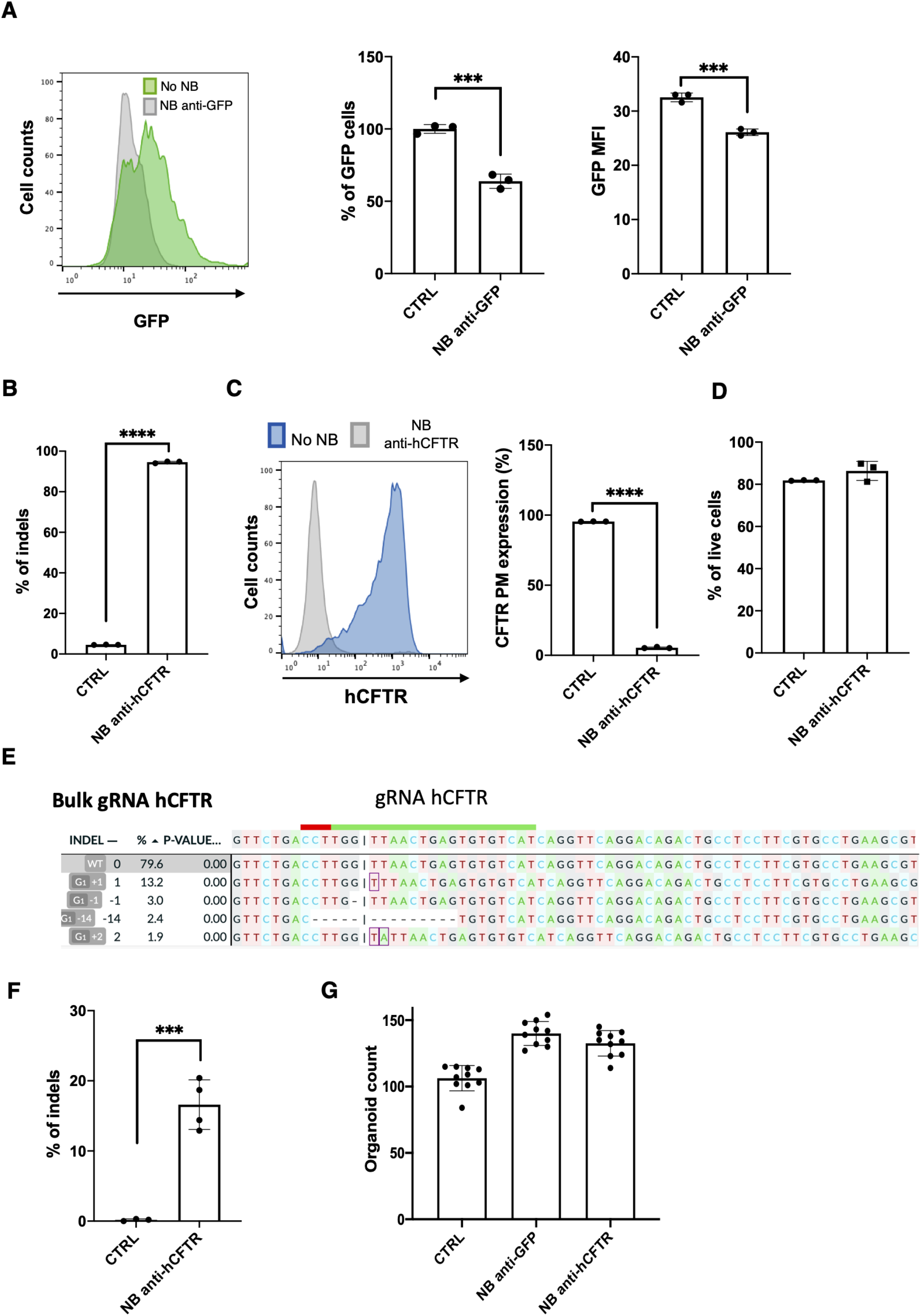
Efficient nanoblade induced knock-out in human intestinal organoids. (**A**) Human rectal eGFP expressing organoids 14 days after treatment or not with anti-GFP nanoblades. Flow cytometry analysis of % eGFP expressing organoid cells (left panels; representative of n=3) and their eGFP mean fluorescence intensity (MFI; right panel) upon treatment with anti-GFP nanoblades (means ± SD, n=3, biological replicates, t-test, Mann Whitney test, non-parametric, ***p<0.001). (**B**,**C**) Validation of anti-hCFTR nanoblades in HEK293T cells. (**B**) One sgRNA was used of KO of human CFTR in HEK293T cells. Bulk gDNA sequencing analysis of the cells treated with nanoblades targeting hCFTR. Sanger sequencing was performed of PCR band including the hCFTR target locus and was subjected to DECODR analysis, which revealed on-target INDEL frequencies at gRNA CFTR target locus (means ± SD, n=3, biological replicates, t-test, Mann Whitney test, non-parametric, ****p<0.0001). (**C**) Detection of hCFTR plasma membrane (PM) expression by FACS analysis for HEK293T cells, overexpressing 3HA-CFTR treated with anti-hCFTR nanoblades or control nanoblades (means ± SD, n=3, biological replicates, t-test, Mann Whitney test, non-parametric, ****p<0.0001). (**D**) Toxicity of nanoblades in HEK293T cells by DAPI staining means ± SD, n=3, biological replicates, unpaired t-test, not significant). (**E**) Bulk gDNA sequencing analysis of human rectal organoids treated with nanoblades targeting hCFTR. Sanger sequencing was performed of the PCR band including the CFTR target locus and was subjected to DECODR analysis which revealed on-target INDEL frequencies at the gRNA hCFTR target locus. Sequence decomposition of INDEL events is shown (upper pannel) and is summarized in the histogram (**F**) (means ± SD n=4, technical replicates, t-test, Mann Whitney test, non-parametric, ***p<0.001). (**G**) Toxicity of nanoblades in organoids is evaluated 14 days after treatment by counting the number of organoids treated or not with nanoblades (means ± SD; n = 10, technical replicates).

Finally, the rectal organoids were upon dissociation incubated solely for 10 min with these validated anti-hCFTR NBs. We amplified a genomic fragment around the loci targeted by the hCFTR sgRNA. The INDEL analysis performed by DECODR software, showed an INDEL frequency up to 20% for the *hCFTR* genomic target site (***Figure 6E&F***). Finally, no apparent toxicity was detected since the number of cells developing into human organoids was similar in the absence or presence of anti-GFP or anti-hCFTR nanoblades (***Figure 6G***).

## DISCUSSION

Here we have shown a novel delivery methodology for the CRISPR/Cas9 system called ‘nanoblades’, which permitted to obtain high levels of gene editing (20%-80%) in human and mouse organoids without the requirement for enrichment via drug selection or fluorescent reporter isolation (Okamoto et al., 2021). These are levels of gene editing that are otherwise exclusively achieved in induced pluripotent stem cells (IPSC) derived organoids (Son et al., 2022). Brain organoids for example are solely derived from iPSCs and since gene editing is performed at the level of the iPSCs, which is highly efficient, deleting genes or introducing genes (e.g. oncogenes) can be performed with ease (Ogawa et al., 2018). In comparison, ASC-derived organoids, showed very low levels of gene editing using the same gene editing tools.

Nanoblades, in contrast to other gene editing techniques, have major advantages: they incorporate the Cas9 associated with gRNA cargo as RNPs which they deliver via pseudotyped VLPs. This allows, in contrast to lentiviral vector delivery (Gu et al., 2022), a transient expression of Cas9. Until very recently (Dawei et al., 2020), most studies performing gene editing in organoids used plasmid DNA to introduce the gene editing machinery. For example, two studies published in 2013 and 2015 introduced plasmid DNA by liposome transfection into adult stem cells (Drost et al., 2015; Schwank et al., 2013). The study published in 2013, which described the use of the CRISPR/Cas9 system to correct the CFTR mutations in organoids derived from cystic fibrosis patients, showed low levels of gene editing (Schwank et al., 2013). The same research team reported the use of CRISPR/Cas9 to induce sequential cancer mutations in cultured human intestinal stem cells; they achieved also in this context low levels of gene editing (< 1%). In these two cases these low levels of gene editing might depend on the delivery method (Drost et al., 2015). Indeed, a study published in 2015 (Fujii et al., 2015), showed higher levels of gene editing. In this case the introduction of DNA plasmid into organoid cells was performed by electroporation. This change of delivery method allowed a fourfold increase in efficiency. However, the levels of gene editing still did not exceed 1% (Menche and Farin, 2021).

To counteract these low efficiency levels, which makes the use of CRISPR/Cas9 in organoids time-consuming and inefficient, several groups developed more efficient reporter gene knock-in techniques (Artegiani et al., 2020; Dawei et al., 2020; Hendriks et al., 2021). The insertion of a fluorescent cassette into the Cas9-induced DSB allows for easier identification of the edited organoids. Despite the advantage of this approach, it should be noted that the addition of a knock-in cassette adds an additional difficulty because a donor template sequence needs to be co-introduced together with the gene editing system into the cell. Furthermore, the knock-in levels achieved by electroporating plasmid DNA remained below 5% (Artegiani et al., 2020).

Viruses have already been used for a long time to genetically manipulate cells. The efficiency of the best performing viral vectors such as lentiviral vectors can reach 100% transduction of the targeted cells (Levine and Friedmann, 1991). In this study, as proof of principle, we evaluated for the first time VLPs incorporating CRISPR/Cas9 anti-GFP gRNA complexes called ‘nanoblades’ for eGFP KO in eGFP expressing human and murine organoids. We reached up to 80% of edited cells, underlining that this technique is more efficient than any other methods inducing a gene KO in organoids.

A crucial concern for any Cas9-mediated gene editing technology is the possibility of off-target editing. When Cas9 cDNA is introduced into cells by a plasmid, it results in a sustained presence of Cas9 protein in the cells and this increases the risk of INDELs at off-target sites. A recent study reported the nucleofection of RNPs to edit organoids (Dawei et al., 2020; Sun et al., 2021). This approach had two advantages. First, in a detailed comparison of the different types of cargo to deliver Cas9, RNPs coupled to single strand gRNAs were more efficient than plasmid DNA encoding Cas9 for editing cells. The second advantage is the reduction of off-target effects. Indeed, by providing Cas9 as a protein, its presence in the cellm is considerably reduced. Therefore, the off-target effects are decreased (Lino et al., 2018). In this study, we provided Cas9 as RNPs delivered by nanoblades. We checked off-target effects of the sgRNAs used, and none of them induced non-specific cleavage at the analyzed off-target genomic sequences that were evaluated by sanger sequencing and ICE and DECODR analysis. Of note, the repair of a DSB induced by CRISPR/Cas9 is error-prone and can give unwanted mutations at the target as well as the off-target genomic loci. A way to avoid these unwanted editing events in organoids might be to use CRISPR-based prime editing (Geurts et al., 2021b) or base editing. Especially, since Liu et colleagues recently showed that they were able to develop nanoblade-like VLPs harboring the base editing machinery and achieve efficient base editing (Banskota et al., 2022).

Another advantage of the nanoblades is the absence of toxicity for the treated organoid cells. Indeed, electroporation remains harsh for the cells. Mammalian cells are sensitive to the voltage and the time of application of the current, even when electroporating RNPs (Lino et al., 2018).

In this study, we reported the generation of an *AR* KO organoid prostate line. As shown previously, luminal cells KO for *AR* failed to achieve terminal differentiation (Xie et al., 2017). Additionally, it was reported that the swelling of luminal organoids is directly dependent on the stimulation of the androgen pathway (Karthaus et al., 2014). In accordance, the organoids KO for *AR* that we generated here induced a compact phenotype. The lumen formed by fluid secretion from the luminal cells did not occur when the cells were KO for *AR*. It would be interesting to understand whether the absence of lumen is indeed due to impaired terminal differentiation of luminal cells. Further studies, using single-cell RNA sequencing for example, would allow to fully understand this observation.

We have also demonstrated for the KO of two different genes (AR and CFTR) that we can multiplex the nanoblades with two gRNA directed against different target sequences of the same gene. Previously, it was demonstrated that nanoblades can be loaded with multiple gRNAs, even with up to 4 different gRNAs (Gutierrez-Guerrero et al., 2021; Mangeot et al., 2019b). Therefore nanoblades might pave the way to study tumorigenesis in gastrointestinal malignancies (Jefremow et al., 2021) and beyond. It is well known that cancers develop through sequential accumulation of oncogenic mutations. Nanoblades will allow generate KOs in multiple genes simultaneously to study their role in cancer development as performed before by fluorescent molecule knock-in for ovarian cancer (Lõhmussaar et al., 2020).

Overall, nanoblades represent a versatile and highly efficient tool to edit human and murine organoids in vitro. They allow to obtain rapid and efficient gene knockout in several organoid models with gene editing levels that outperform other currently used techniques, while requiring only transient expression of the gene editing machinery. Of utmost importance, efficient gene editing in organoids in accompanied by low cellular toxicity and low off-target effects. Finally, the experimental strategy is fast since it allows to generate organoid lines in 4 to 6 weeks with high efficiency. Finally, nanoblades offer the possibility to efficiently generate high level editing in organoids without the need to use reporter encoding knock-in cassettes or a drug selection method. Therefore, nanoblades might allow studies in tissue differentiation, cancer development and drug screening, as well as preclinical evaluation of correction by gene therapy in organoids and possibly will allow to facilitate multiple other gene editing applications using organoids.

## Acknowledgements

Funding was received from the CHEMAAV-ANR-19-CE18-0001 grant operated by the French National Research Agency (ANR). This study was supported by research funding (CRISPR screen Action) from the Canceropôle Provence-Alpes-côte d’Azur, the French National Cancer Institute (INCa) and the Provence-Alpes-côte d’Azur Region. AK and VT received a PhD fellowship from the French Ministry of Research. M.B. and M.M.E. are supported by an FWO-SB (Flemish Research Foundation) doctoral fellowship 1SE8122N and 1S29917N respectively; M.S.C. by a senior post-doctoral FWO scholarship 12Z5920N and KU Leuven BOFZAP professorship. Cystic fibrosis research is funded by the Belgian CF patient Association and Fund Alphonse Jean Forton from the King Baudouin Foundation (2020-J1810150-E015) and by FWO (SBO OrganID; S001221N).

## Author contributions

VT and AK were responsible for experimental design, methodology, experiments, analysis, and preparation of the first draft of the manuscript. EB, MB, MME, MHG, DH, FL, AG, LM were responsible for designing, performing and analyzing experiments. FV and PEM provided materials and advise. RG, MSC, HC and were involved in design and supervision of the human organoid experiments, software analysis, and in the review and editing of the paper. FB was involved design, funding acquisition, supervision of the murine organoid experiments in the review and editing of the paper. EV designed the project, supervised the work, performed experiments, discussed and analyzed results and wrote the manuscript.

## Data availability statement

The source data including sanger sequencing data supporting the conclusions of this article are made available.

## Supplementary Files

Supplemental Figure 1

Supplementary Tables 1, 2, 3, 4 and 5

---

**Supplemental Figure 1.**
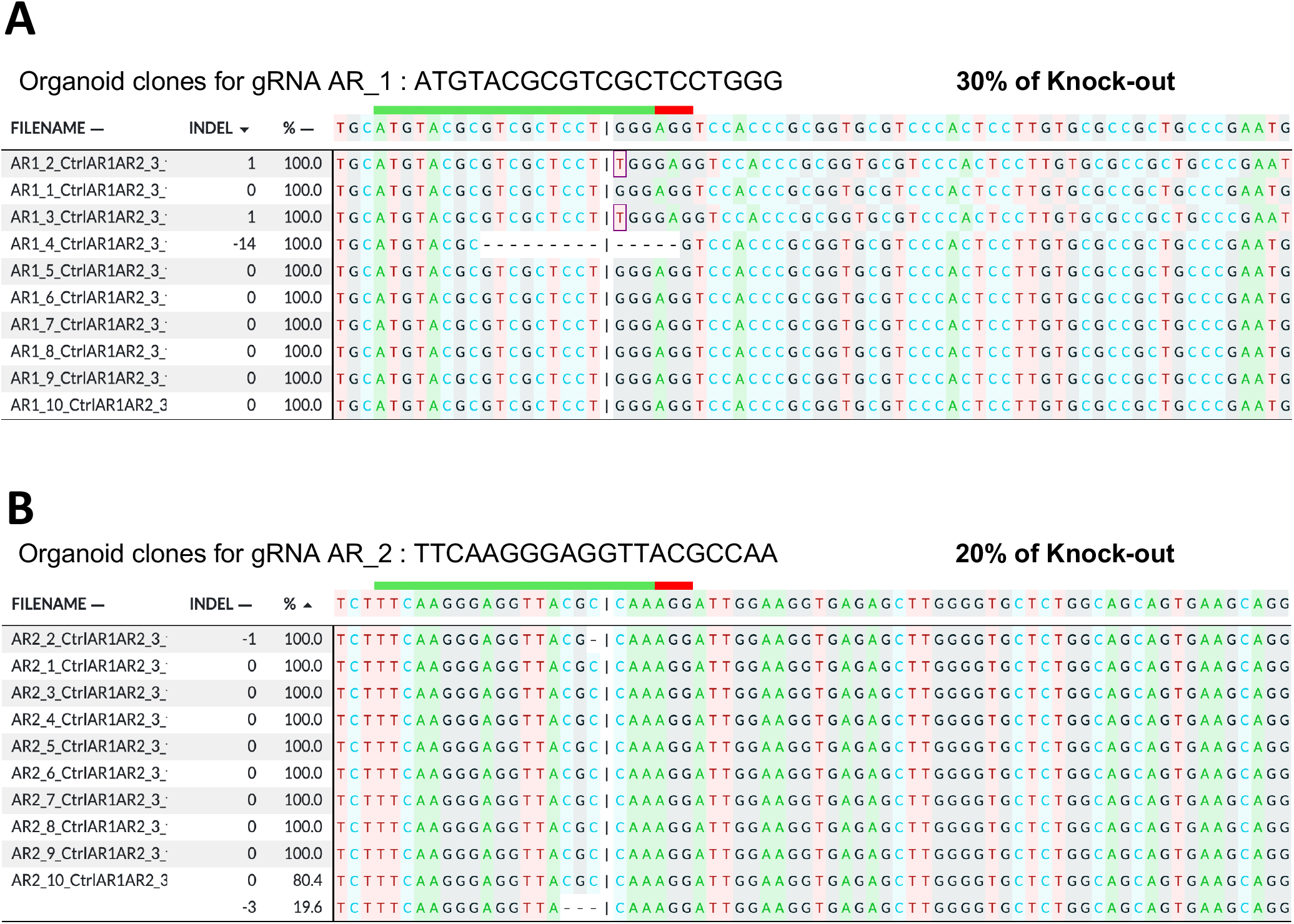
Nanoblades enable efficient generation of knock-out organoids clones from mouse prostate organoids. Mouse prostate organoids at 9 days after treatment with nanoblades targeting the Androgen-Receptor (AR). Two different sgRNAs (AR_1 and AR_2) were used to knock-out AR. Single organoids were harvested, gDNA was isolated and sequenced for the loci targeted by the different AR guide RNAs. Sanger sequencing was performed of the 557 bp PCR surrounding the AR target loci and was subjected to ICE analysis which revealed on-target INDEL frequencies at gRNA AR1 (**A**) or AR2 (**B**) target loci.

